# Intense Pulsed Electric Fields Denature Urease Proteins

**DOI:** 10.1101/572784

**Authors:** Gen Urabe, Toshiaki Katagiri, Sunao Katsuki

## Abstract

This paper describes the effects of nanosecond pulsed electric fields (nsPEFs) on the structure and enzyme activity of three kinds of proteins. Intense (up to 300 kV/cm), 5-ns-long electrical pulses were applied to solutions of lysozyme (14 kDa, monomer), albumin (67 kDa, monomer), and urease (480 kDa, hexamer). We analyzed the tertiary and quaternary structures of these proteins as well as their enzyme activity. The results indicated the deformation of both the quaternary and tertiary structures of urease upon exposure to an electric field of 250 kV/cm or more, whereas no structural changes were observed in lysozyme or albumin, even at 300 kV/cm. The enzyme activity of urease also decreased at field strengths of 250 kV/cm or more. Our experiments demonstrated that intense nsPEFs physically affect the conformation and function of some kinds of proteins. Such intense electric fields often occur on cell membranes when these are exposed to a moderate pulsed electric field.

## Text

Electroporation (or electropermeabilization) has been widely confirmed and several attempts to investigate this phenomenon using lipid vesicles have suggested that electric pulses can damage plasma membranes and lead to their disruption(1). Numerous studies have derived from this insight. Electrochemotherapy, Ca electroporation, and tumor ablations for medical applications; non-thermal pasteurization; and electroextraction for food-processing applications based on nanosecond pulsed electric fields (nsPEFs) are attractive techniques and there have been many attempts to put them into practice(2, 3). There are numerous accounts describing that nsPEFs lead to several biological responses, whose kinetics have been partially obvious, such as cell-morphology transformations, stress responses, signal transductions relevant to cell death, and calcium-ion reactions(4, 5). Most of these responses have been attributed to an abrupt increase in the concentration of calcium ions, which work as messengers to signal diverse biological reactions, triggered by the permeabilization of the plasma membrane or the surface of the endoplasmic reticulum, promoting trans-membrane calcium mobilization into the cytoplasm(6, 7).

Other cell components such as proteins, which are electrically charged dielectric compounds, receive stress from the fields. In particular, membrane proteins are exposed to extremely high electric fields in the order of MV/cm because the electric field is enhanced on the membrane, which is a sub-10-nm-thick dielectric film, owing to polarization and charge accumulation under an external field. Several numerical calculations predict that proteins could respond to electricity. Microtubules are able to transmit electrical pulses along their structures and have a dipole moment, which generates electric fields around them(8–11). However, there are only few reports on experimentally proven electrical effects on proteins. Amino-acid residues of a crystallized protein were found to change their direction when exposed to a 1 MV/cm electric field(12), but because a protein crystal was used instead of a solution, the conditions did not represent the physiological ones. Therefore, it is important to evaluate the effects of nsPEFs on proteins under physiological conditions to understand their physical impact and the subsequent biological reactions.

In addition, membrane proteins are under substantial electric fields in the order of 100 kV/cm because of the residual membrane potential of approximately 70 mV. It may be necessary to take electrical perspectives into consideration to completely understand the biology. Several papers suggest that electric fields affect the cell activity, and most of the proposed mechanisms are supported by numerical calculations^13–15^.

There are several applications of studies on proteins exposed to high electric fields; controlling intracellular electric fields may enable the manipulation of cell activities such as cell division, which could suppress tumor growth(15, 16). A previous study reported that a bacterial spore wall composed of peptidoglycan was damaged by intense pulsed electric fields of 7.5 kV/cm(17). It is critical to examine electrical impacts on proteins from the viewpoints of bioelectrics, basic biology, and applications.

Here we focused on proteins in a liquid to simulate the intracellular conditions and analyzed irreversible responses. Because proteins have four hierarchical structures—from primary to quaternary—we were interested in studying the destructive effect of intense electric fields on the class of structure, molecular size, and chemical binding. This study discusses the deformations of the primary, tertiary, and quaternary structures using several proteins with different structural features. We have developed a 5-ns-long high-voltage pulse generator to apply intense electric fields to protein solutions in a 1-mm-gap cuvette. Voltage pulses such as the one shown in Fig. 1, measured at the cuvette using a capacitive divider, were repetitively delivered to the cuvette at a repetition frequency of 3 Hz(18). The maximum electric-field strength in the cuvette was 300 kV/cm, and repetitive pulsing had a small thermal effect with a temperature rise of 3°C.

**Figure. 1.**
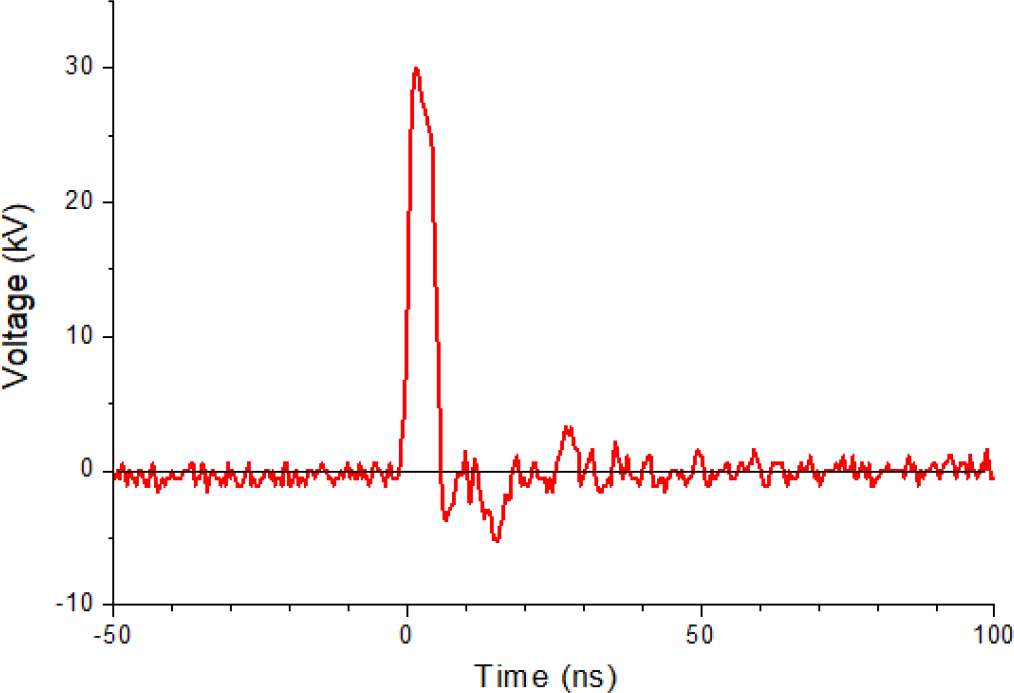
Typical waveform of the voltage applied to the 1-mm-gap cuvette containing the protein solution.

We purchased three proteins with various structures and molar weights (all from Wako Chemicals), namely, lysozyme from egg white (14 kDa, monomer), albumin from bovine serum (67 kDa, monomer), and urease from jack bean (480 kDa, hexamer), and dissolved each protein into D-PBS to obtain 1 mg/ml protein solutions.

To analyze the effects of an electric field on the primary structure of the proteins, we exposed lysozyme and albumin to nsPEFs and examined their structures using SDS-PAGE. The bands observed for all the samples were the same as those obtained without PEF exposure, suggesting that the electric fields were unable to cut the peptide bonds [Figs. 2(A), (B)].

**Figure. 2.**
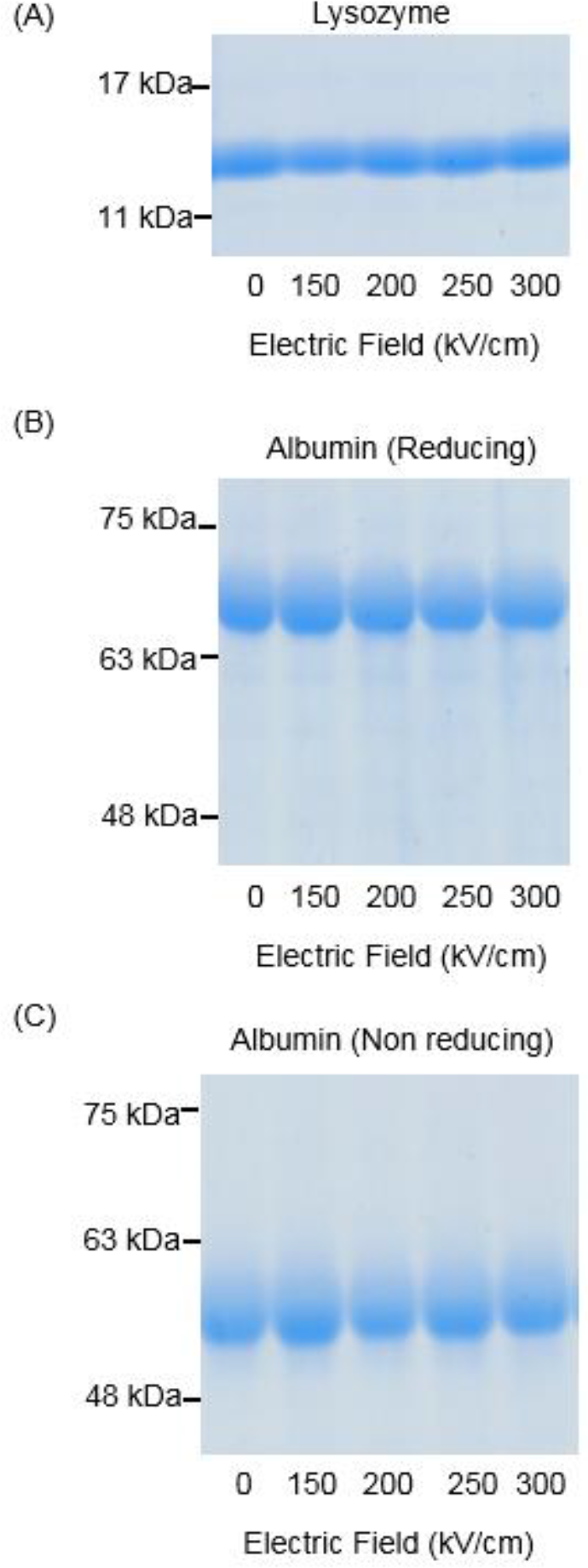
SDS-PAGE of lysozyme and albumin exposed to nsPEFs: (A) Electrophoresis of lysozyme mixed with a reducing agent. The numbers below the gels represent the strength of the nsPEF. The figures on the left side of the gels show the positions of the markers with the written molar weights. (B,C) Electrophoresis of albumin with (B) and without (C) reducing agent.

The tertiary structures involve two binding types, namely, disulfide bonds and non-covalent bonds. To evaluate the damages on the disulfide bonds, we compared albumins under two different treatment conditions using SDS-PAGE: one of the arrangements left the disulfide bonds intact (non-reducing conditions), whereas the other one broke them (reducing conditions). The two conditions differed in whether 2-mercaptoethanol (2ME) (reducing agent) was added to the samples. Albumin’s band pattern did not change under reducing or under non-reducing conditions, which implies that a 300 kV/cm nsPEF does not break the disulfide bonds [Fig. 2 (B), (C)].

Urease, which is free from covalent bonds in the tertiary and quaternary structures, was chosen as a large “soft” protein. In the normal SDS-PAGE protocol, proteins are preheated at 95°C for 5 min with SDS(surfactant) and 2ME to destroy the tertiary and secondary structures with digesting disulfide bonds and non-covalent bindings. It is impossible for normal protocol to investigate the tertiary and quaternary structures. In this study, we lowered the preheating temperature and shortened the time to prevent the high-order structures in urease from being completely destroyed. Preheating at 60°C for 2 min was chosen to keep the bands of the trimer and subunit emerging simultaneously with the same intensity. Although the main structure of urease is a hexamer, only the trimer and subunit bands were visible because the hexamer is too large for the electrophoresis performed in this study [Fig. 3(A)]. Under non-reducing condition, the trimer bands shifted downward and smears appeared below the subunit bands when the applied electric fields were 250 kV/cm or more [Fig. 3(A), (B), (C)]. The trimer band also shifted under reducing conditions [Fig. 3(A), (D)], which implies that new disulfide bonds could be produced between the cysteine residues in the proteins upon exposure to nsPEFs(19). This can be a reasonable explanation for the fact that smears were observed below the subunit under non-reducing conditions but disappeared under reducing conditions. The urease subunits might be able to form new disulfide bonds because they contain cysteines without disulfide bonds. However, the trimer bands of urease also jumped under the reducing conditions, suggesting that nsPEFs not only produce disulfide bonds but also have other effects on the proteins. It has been reported that electric fields can damage hydrogen bonds, thereby changing the secondary structures(20). In agreement with a previous study, the urease trimer bands shifted downward under both reducing and non-reducing conditions.

**Figure. 3.**
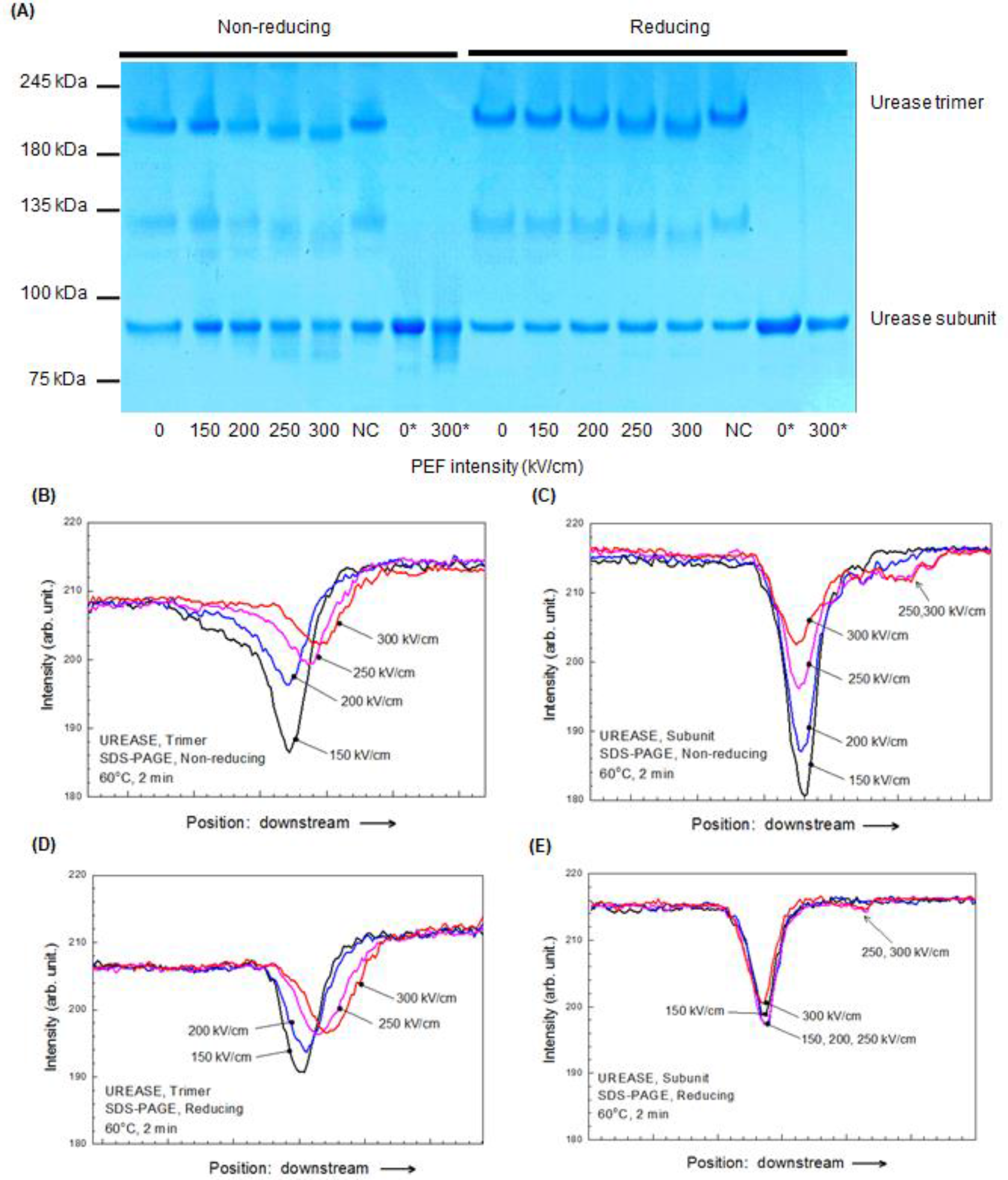
(A) SDS-PAGE of urease treated at 60°C for 2 min under non-reducing or reducing conditions. The bands between 180 and 245 kDa correspond to the trimer where as those between 75 and 100 kDa correspond to the subunits. The numbers below the gel represent the strengths of the nsPEF. NC is the sample in which normal urease is dissolved in nsPEF-treated PBS. 0^*^ and 300^*^ are the samples that were exposed to nsPEF strengths of 0 and 300 kV/cm, respectively, and boiled at 95°C for 5 min. (B–E) Band-intensity distributions of SDS-PAGE: trimer bands under non-reducing conditions (B), subunit bands under non-reducing conditions (C), trimer bands under reducing conditions (D), and subunit bands under reducing conditions (E).

Although Fig. 3(A) suggests that the sum of the trimer and subunit amounts decreases with increasing electric-field strength, we do not discuss the band-intensity variations because the hexamer was not shown under the present SDS-PAGE conditions.

Protein structures are important and determine their functions. We analyzed the enzyme activity of urease exposed to nsPEFs using the QuantiChrom Urease Assay Kit (BAS, DURE-100). The enzyme activity is determined by the amount of ammonia (in μmol) synthesized in 1 min in 1 L of a liquid. At electric-field strengths of 250 kV/cm or more, the activity significantly decreased [Fig. 4]. This trend coincides with that of the structure changes determined by SDS-PAGE [Fig. 3 (A)].

**Figure. 4.**
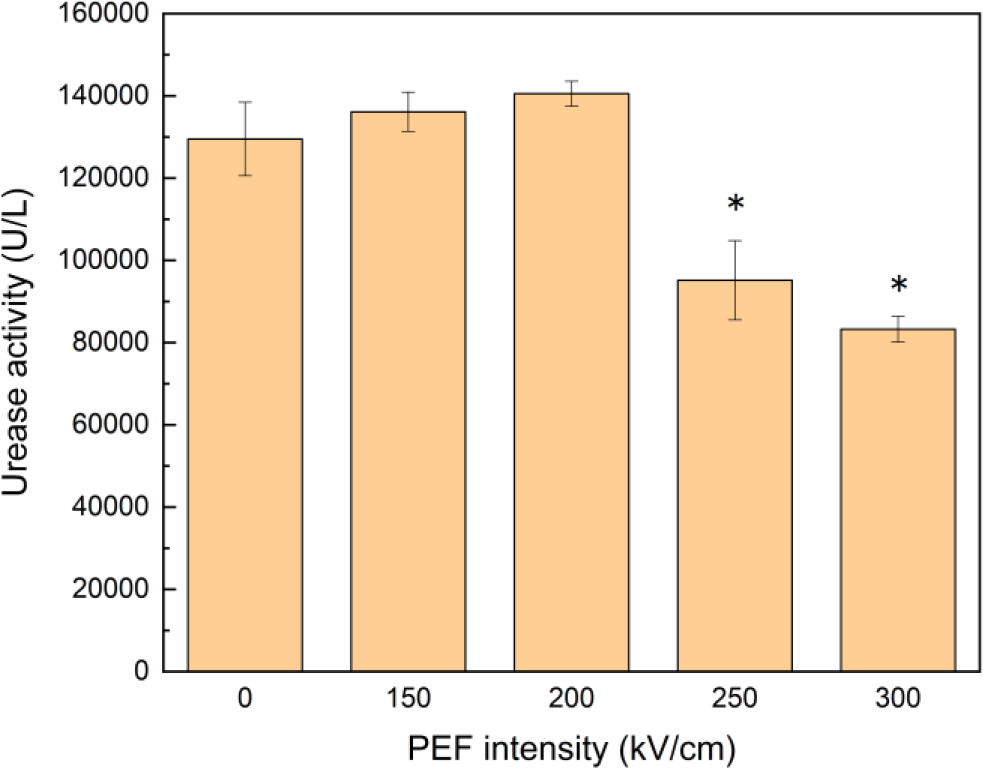
The enzyme activity of urease exposed to nsPEFs with several electric-field strengths. The error bars are described with standard error. *: p < 0.01.

In conclusion, nsPEFs with strengths above 250 kV/cm can affect not only the structure but also the enzyme function of urease, which serves as a representative of relatively large and soft proteins, whereas rigid proteins such as lysozyme and albumin, which have covalent bonds in their high-order structures, seem to remain unchanged upon exposure to nsPEFs up to 300 kV/cm.

## Author Contributions

Sunao Katsuki designed the research. Toshiaki Katagiri built and operated the pulse generator and measured the voltage. Gen Urabe analyzed the proteins and wrote the manuscript.

## Acknowledgement

This study was in part supported by Grant-in-Aid for Scientific Research (17H03220).

